# Distinct modes of action of IAPP oligomers on membranes

**DOI:** 10.1101/2021.06.24.449712

**Authors:** Aliasghar Sepehri, Binod Nepal, Themis Lazaridis

## Abstract

Islet Amyloid Polypeptide (IAPP, also known as amylin) is a peptide hormone which is co-secreted with insulin by pancreatic β-cells and forms amyloid aggregates in type II diabetes. Various lines of evidence indicate that oligomers of this peptide may induce toxicity by disrupting or forming pores in cell membranes but the structures of these pores are unknown. Here we create models of pores for both helical and β-structured peptides using implicit membrane modeling and test their stability using multimicrosecond all-atom simulations. We find that the helical peptides behave similarly to antimicrobial peptides; they remain stably inserted in a highly tilted or partially unfolded configuration creating a narrow water channel. Parallel helix orientation creates a somewhat larger pore. An octameric β barrel of parallel β-hairpins is highly stable in the membrane, whereas the corresponding barrel made of antiparallel hairpins is not. We propose that certain experiments probe the helical pore state while others probe the β-structured pore state; this provides a possible explanation for lack of correlation that is sometimes observed between in vivo toxicity and in vitro liposome permeabilization experiments.

## INTRODUCTION

Type II (non-insulin dependent) diabetes is characterized by a β-cell deficit in the pancreas, increased β-cell apoptosis, and amyloid fibrils of a 37-residue peptide hormone named amylin or Islet Amyloid Polypeptide (IAPP), which is co-secreted with insulin ^1^. The role of the IAPP aggregates as a causative agent of this disease has been scrutinized in the past two decades ^2–4^. While some researchers found fibrils to be toxic to β-cells ^5–8^, others found higher toxicity in smaller oligomeric precursors ^9–13^. A recent study found that the toxic species is soluble oligomers with less than 6 protomers ^14^. Various mechanisms of IAPP toxicity have been proposed, the primary of which is membrane permeabilization ^15^, by pore formation ^10,16–19^, lipid extraction ^20,21^, or fibril growth ^22^. Compromise of the integrity of either the plasma or the ER membranes could lead to high Ca^2+^ concentration in the cytoplasm, triggering apoptosis ^23^. However, it has also been found that the correlation between toxicity and *in vitro* membrane permeabilization is not perfect ^24^. Similar considerations and findings apply to other aggregating proteins and the corresponding diseases, such as amyloid β in Alzheimer’s ^25^ and α-synuclein in Parkinson’s ^26^.

The structure of fibrils of IAPP and other amyloidogenic molecules is dominated by β-sheets ^27,28^ but the details differ. Some earlier studies suggested a U-shaped ribbon ^29^ while others an S-shaped molecule (Kajava et al. 2005). Recent CryoEM structures found S-shaped or wavy patterns ^32–34^. Much less is known about the structure of the toxic soluble oligomers. One study found that they have partial helicity and less than 15% β character ^14^, while another found that the toxic species is oligomers with significant β content ^11^. Crystallography revealed an out-of-register β structure for one 7-residue IAPP fragment ^35^. Out-of-register β structures have been suggested to be toxic in other peptides ^36^. Rawat et al. found small oligomers to be largely helical and large ones β ^37^. A 3-strand β-sheet intermediate was identified ^38^, but its role in toxicity is unclear.

There is considerable evidence that prefibrillar oligomers permeabilize membranes by forming pores or defects ^9,10,39–45^. Electrophysiology characterized the ion channel characteristics of human IAPP, while rat IAPP did not form ion channels ^39^. If oligomers are dominated by β structure, any pores that the peptide makes are expected to also be β-structured. On the other hand, membrane permeabilization has been observed under conditions where the peptide should be a monomeric helix ^16,46^.

Most modeling work on IAPP has been concerned with aggregation in solution ^47,48^. A few articles reported studies of the interaction with membranes and possible pore formation mechanisms ^19,49–51^. Use of the Martini coarse-grained force field to model the peptides as helices inserted in the membrane ^52,53^ revealed possible pore-like structure formed by 5mers or 6mers. Other work investigated the interaction of fibril-like β structures with membranes ^54,55^.

One key piece of information for modeling is the secondary structure of the pore-forming species. Monomeric IAPP is helical in membrane environments ^56–58^ and has a size similar to that of antimicrobial peptides (AMPs). Membrane permeabilization has been observed under conditions where the peptide should be a monomeric helix ^16^. One can envision a number of helices coming together to form a pore, as assumed in the above coarse-grained studies ^52,53^. It has been suggested that the 19 N-terminal residues, which are predominantly helical in the membrane, are responsible for membrane poration ^46,59^. On the other hand, the oligomers thought to cause membrane damage have a mostly β character, with its core in segment 20-29 ^11,14^.

Based on the experimental results summarized above, we hypothesized that there are two distinct mechanisms of pore formation by IAPP: a) helical monomers approaching each other, inserting, and stabilizing a pore, b) oligomers rich in β structure of the appropriate type inserting and forming a β-barrel. The latter is relevant for the toxicity of soluble oligomers. Here we test these hypotheses by implicit-solvent and extensive all-atom simulations. For helical peptides we find structures similar to those in our previous work on AMPs ^60^. An octameric β barrel with parallel β hairpins is found to be highly stable, while a similar barrel with antiparallel hairpins is not. We discuss these findings in the context of known experimental facts and consider possible ways of experimental validation of these structures.

## METHODS

### Implicit solvent modeling

Implicit membrane simulations employed the IMM1 model^61^, an extension of the EEF1 effective energy function for soluble proteins^62^ to heterogeneous membrane-water systems. IMM1 uses a switching function that transitions smoothly from a nonpolar to an aqueous environment. It accounts for the surface potential using the Gouy-Chapman theory^63^. Modeling of pores^64,65^ is accomplished by making the switching function F dependent not only on the vertical (z) coordinate but also on the radial coordinate (the distance r from the z axis):

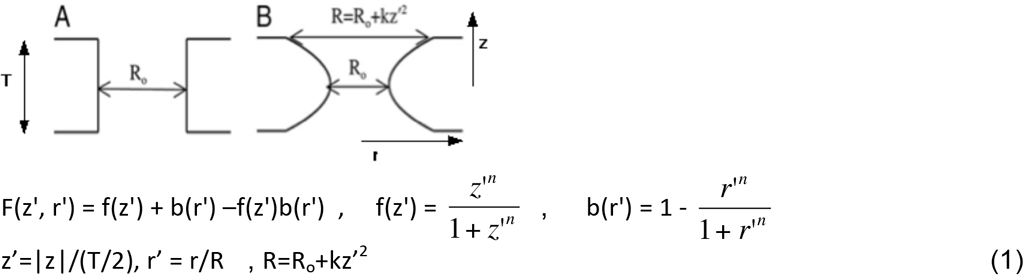

The sequence of IAPP is KCNTATCATQ R**L**AN**FLV**HSS NN**F**GA**IL**SST N**V**GSNT**Y**-NH2, with a 2-7 disulfide bond (in bold are the hydrophobic residues). The peptide was built as an α-helix at residues 8-38 with a disulfide bond between 2 and 7. It was then placed on a flat implicit membrane with its hydrophobic side facing the membrane or into a cylindrical pore in a transmembrane orientation, also with its hydrophobic side facing toward the membrane.

To consider the possibility of β barrel formation, the sequence was scanned for triplets of nonpolar residues that would face in the same direction in a β strand conformation (i.e. ΦxΦxΦ, where Φ is a hydrophobe). This is based on the expectation that stability of a β strand in the membrane requires the burial of ideally 3 hydrophobic side chains. One of them need not be strongly hydrophobic (e.g. A or G) but it cannot be strongly polar. Not many possibilities exist in the IAPP sequence. One AxFxV triplet exists at 13-17 and one FxAxL triplet at 23-27. One or both of them could contribute to an oligomeric β barrel. Here we consider the latter possibility, i.e. each peptide contributes 2 strands, one at 13-17 and another at 23-27. We constructed an octameric β barrel using our previous β barrel of 8 protegrin β-hairpins ^66^ as a template based on the following alignment:

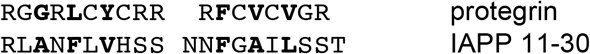

First, a single hairpin of IAPP was generated by mutating a single protegrin 1 β hairpin from the template. The side chains of IAPP and the missing N-terminal and C-terminal residues were built in an extended conformation. The parallel IAPP barrel was then generated by replicating and translating a single IAPP hairpin along a circle of radius 15 Å at multiples of a 45° angle. The tilt angle of each IAPP strip with the barrel axis was 38°. The resulting structure was then subjected to a 5-ns unrestrained MD simulation using IMM1 in a neutral toroidal pore of radius R_o_=15 Å and curvature parameter k=17 Å (see Eq. 1).

An antiparallel barrel was constructed in a similar way but here every other hairpin was turned upside down by rotating 180° around an axis perpendicular to the hairpin crossing its center of mass. This way the same residues face inside as in the parallel barrel. The constructed barrel was then subjected to a 5-ns MD simulation in an IMM1 neutral toroidal pore (R_o_= 15 Å, k=17 Å). However, here a cylindrical restraint using the MMFP module in CHARMM was found necessary to exclude the hairpins from a 9 Å radius.

Binding energies to the membrane were estimated as average effective energies of transfer (ΔW) of the membrane-embedded configurations from water to the membrane.

### All-atom simulations

For the helical bundle simulations, we applied the approach we followed previously for AMPs^60,67^. Six peptides were placed on a circle of radius ~6 Å along the z-axis (perpendicular to the membrane) in parallel and antiparallel orientations with the hydrophobic face oriented toward the lipids and hydrophilic residues toward the pore. The hexamer was uploaded onto the CHARMM-GUI server^68^ where 180 lipids (POPC: POPG 7:3), a water slab at least 17.5 Å thick, and neutralizing ions (K^+^) were added to the system. The parallel and antiparallel barrels from the IMM1 simulations were treated similarly. Potassium chloride (0.15 M) was added with extra K^+^ to neutralize the excess charges.

The four systems were equilibrated using the NAMD software package^69^ in six steps, each run for 25 ps with 1-fs time step. In the first equilibration step, harmonic constraints (k = 1 kcal/mol/Å^2^) were applied to the water atoms, ions, phosphorous atoms, and peptide backbone atoms. The constraints on the lipids, waters and ions were subsequently released one at a time in each subsequent step. The last step was an unconstrained equilibration with a time step of 1 fs followed by 5 ns run with a time step of 2 fs. The final structures were subjected to ANTON2 simulations. Table 1 summarizes the properties of these four systems.

**Table 1.** Systems subjected to all-atom simulations, each for 10 μs except for the antiparallel β barrel which was run for 1 μs

| System | Peptides | Atoms | Water | Lipids | POPC Upper | POPC Lower | POPG Upper | POPG Lower | Ion |
| --- | --- | --- | --- | --- | --- | --- | --- | --- | --- |
| IAPP Helix Parallel | 6 | 59400 | 10800 | 180 | 64 | 62 | 26 | 28 | 36 $K^+$ |
| IAPP Helix Antiparallel | 6 | 61089 | 11363 | 180 | 63 | 63 | 27 | 27 | 36 $K^+$ |
| IAPP Beta barrel Parallel | 8 | 73461 | 14225 | 200 | 67 | 73 | 30 | 30 | 73 $K^+$<br>37 $Cl^-$ |
| IAPP Beta barrel Antiparallel | 8 | 80088 | 16430 | 200 | 69 | 71 | 30 | 30 | 79 $K^+$<br>43 $Cl^-$ |

## RESULTS

### Implicit solvent modeling

Except for the 7 N-terminal residues linked by a disulfide bond, IAPP can form an amphipathic helix. Three hydroxyl residues (S19, S20, T30) on the hydrophobic face compromise its hydrophobicity and make binding to a zwitterionic membrane weak (α-synuclein shares this characteristic ^70^). Weak binding was also seen in all-atom free energy simulations ^19^. Consistent with experiment ^71–75^, the peptide binds more stably to a 30% anionic membrane due to the favorable interaction of Lys, Arg and the N-terminus with the membrane anionic charge (ΔW = −6.6 ± 0.8 kcal/mol, Fig. 1a). The helicity at the C-terminus, well maintained in up to 10 ns MD simulation, is probably overestimated, since EPR studies found a helix only at residues 9 to 22 on 80% POPS vesicles ^56^. The IAPP helix also binds favorably to implicit membrane pores. The binding energy to a 13-Å radius toroidal pore is −3.8 ± 0.8 kcal/mol and to a cylindrical pore −4.9 ± 1.0 kcal/mol (Fig. 1b). Thus, helical IAPP is plausible as a helical pore-forming peptide.

**Fig. 1.**
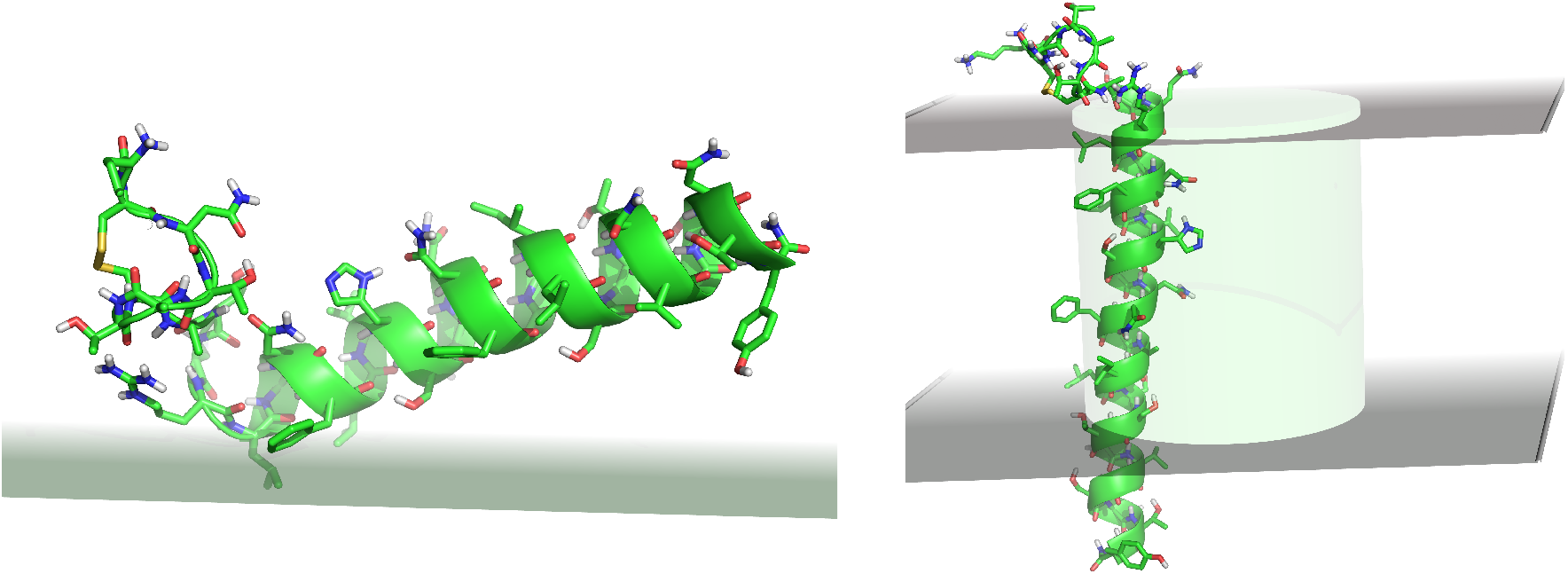
IAPP on the surface of a 30% anionic implicit membrane (left) and a 13-Å radius implicit cylindrical pore (right). The grey planes are the boundaries of the membrane’s hydrophobic core.

To examine the plausibility of a β-barrel pore, we constructed one at residues 11-30 using a previous protegrin barrel as a template (see Methods). This model (Fig. 2) in an implicit toroidal pore (Ro=15, k=17) is stable upon MD simulation and gives a large, favorable transfer energy from water to the pore (−35 ± 2 kcal/mol). Most N-terminal segments extend outward from the membrane, with occasional intrusions into the pore lumen. The model is not stable in a cylindrical pore of the same radius and partially moves out. This binding energy to the pore is reasonable compared to other β barrel proteins. For example, OmpA (pdb id 1bxw) gives −16 kcal/mol and a recently designed β barrel (pdb id 6×9z ^76^) gives −25 kcal/mol (both in R=9 Å cylindrical pores). A barrel made of antiparallel β-hairpins required a cylindrical exclusion restraint to maintain its structure, otherwise it collapsed to a β-sandwich. The reason seems to be the inability of this structure to keep all polar and charged residues out of the nonpolar membrane environment. Favorable interactions between alternating N and C termini in the parallel barrel may also contribute to its higher stability.

**Fig. 2.**
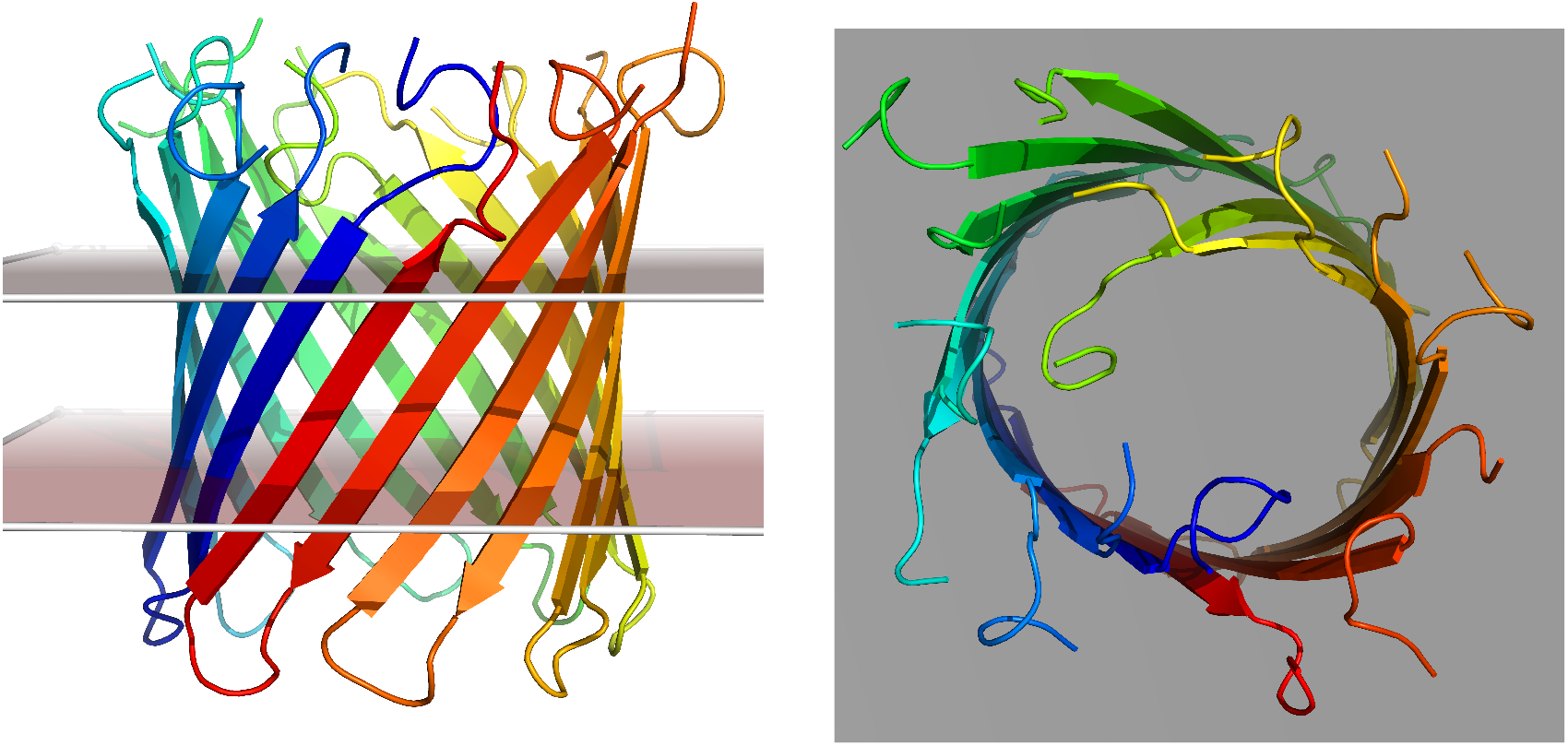
IAPP octameric parallel barrel in an implicit toroidal pore. Side view (left) and top view (right). The grey planes are the boundaries of the membrane’s hydrophobic core.

It is interesting to compare the energy of the peptides in the various possible states: monomeric helix on the membrane, β hairpin in the parallel barrel, and β strand in the fibril. To that end, one of the recent fibril structures was considered ^33^. The 12 peptides from the PDB file were simulated in implicit water after addition of the missing N-terminal residues 1-12 in extended conformation (with the disulfide bond). The effective energy of a peptide in an infinite fibril was calculated as the intramolecular energy plus one half of the interaction energy with its surroundings. The resulting values are shown in Table 2. These values include only peptide-peptide energies and solvation free energies; they do not include peptide configurational entropy, which would be more favorable for the monomeric helix, and membrane deformation free energy for the barrel. These caveats notwithstanding, the energies in Table 2 are consistent with the common notion that the fibril is the global free energy minimum. The barrel is more stable than the monomeric helix, but metastable with respect to the fibril.

**Table 2.**
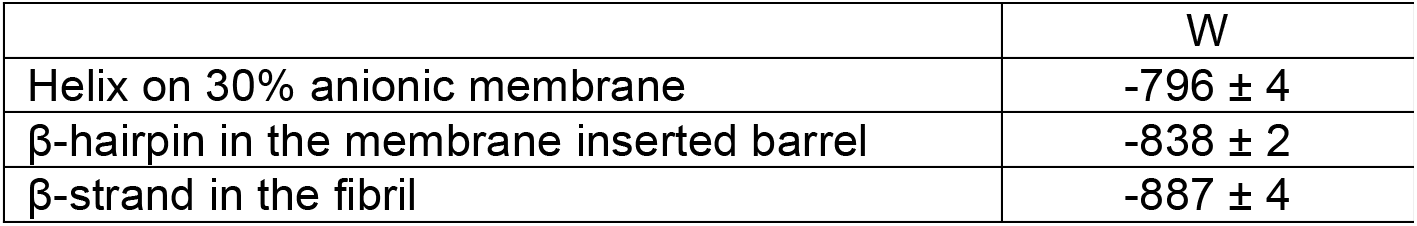
Effective energy of a monomer (kcal/mol) in three states

### All-atom simulations

In our previous work on melittin-derived pore-forming peptides^67^, we observed that a large pore occurs where the attractive interactions between adjacent peptides are stronger than those between non-adjacent ones. We simulated parallel helices and saw that non-adjacent interactions happen between unfolded residues close to C-terminus. We also found that interactions that can keep adjacent peptides tilted towards each other, such as interactions between a residue close to the C-terminus and a residue close to the middle of the peptide, cause a large pore.

The initial configurations of the IAPP parallel and antiparallel helix bundles consist of six long helices with 30 residues, i.e. residues 8-37, perpendicular to the lipid bilayer (Figs. 3a and 4a). This is longer than melittin-like peptides with 26 residues. So, in order to maintain helicity, a peptide needs to be highly tilted; this occurs for peptides A, D and F in parallel configuration and peptides A and F in antiparallel configuration. Other peptides remain close to the inserted state (low tilt angle) by unfolding some residues near the C-terminus that are located in the aqueous phase. Helical remain the regions 8-31, 8-29, and 11-27 for peptides E, B, and C, respectively, in the parallel bundle, and regions 8-28, 8-31, 8-34, and 8-30 for peptides B, C, D, and E, respectively, in the antiparallel bundle. In both systems, the tilt angle and helicity of each peptide do not significantly change after 2 μs and most of changes in peptide configuration happen at the termini (Figs. 3b, 3c, 4b and 4c). The antiparallel system is organized in the form of three antiparallel dimers, one of which is less tight than the others.

**Fig. 3.**
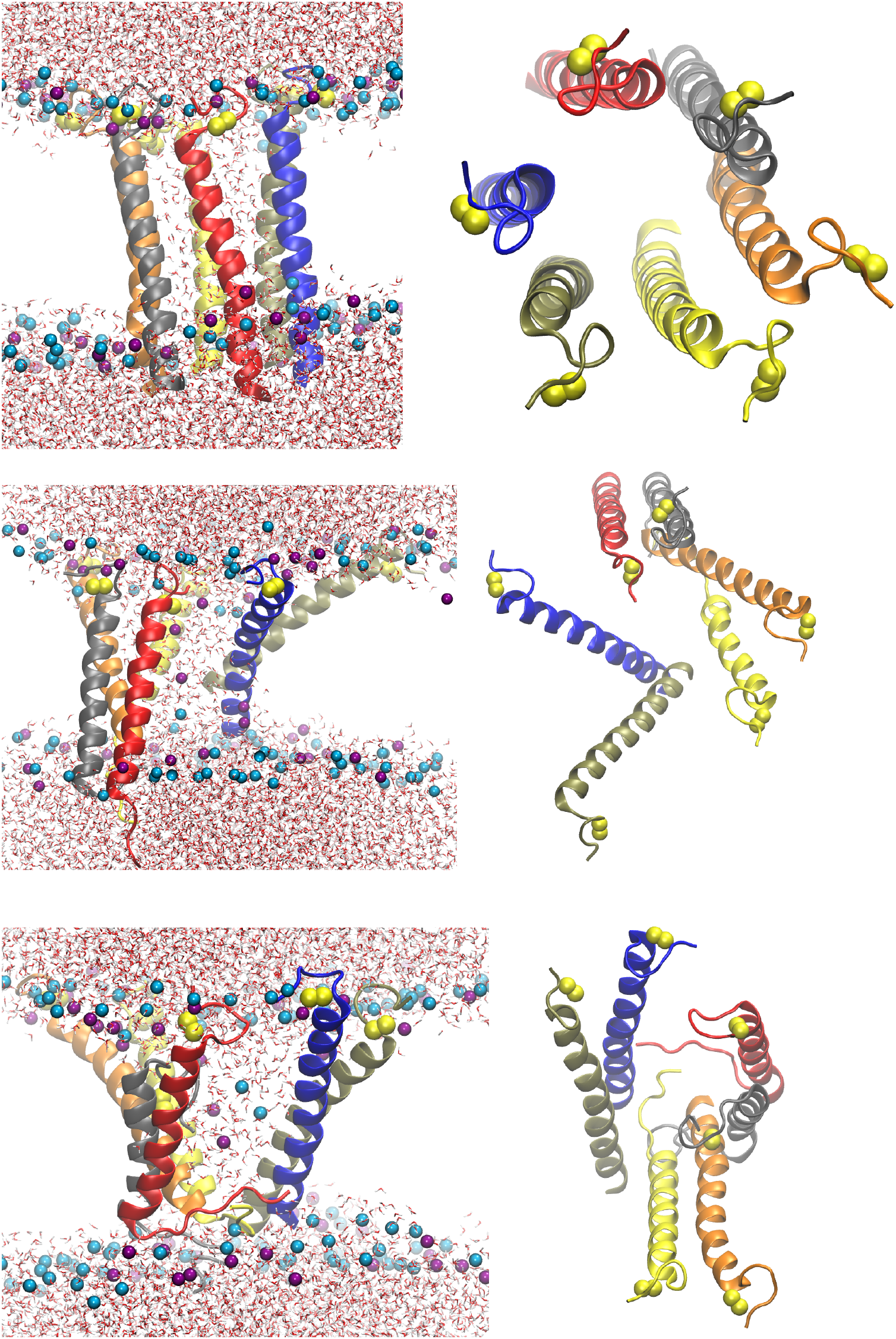
Parallel IAPP helices in initial configuration (above), after 2 μs of the simulation (middle), and at the end of the simulation (10 μs, lower). Left panels side views, right panels top views. Color code: POPC headgroup = cyan blue spheres, POPG headgroup = purple spheres, Sulfur atoms = Yellow spheres. Peptides: A = blue, B = red, C = charcoal, D = orange, E = yellow, F = olive, G = light grey, H = green.

**Fig. 4.**
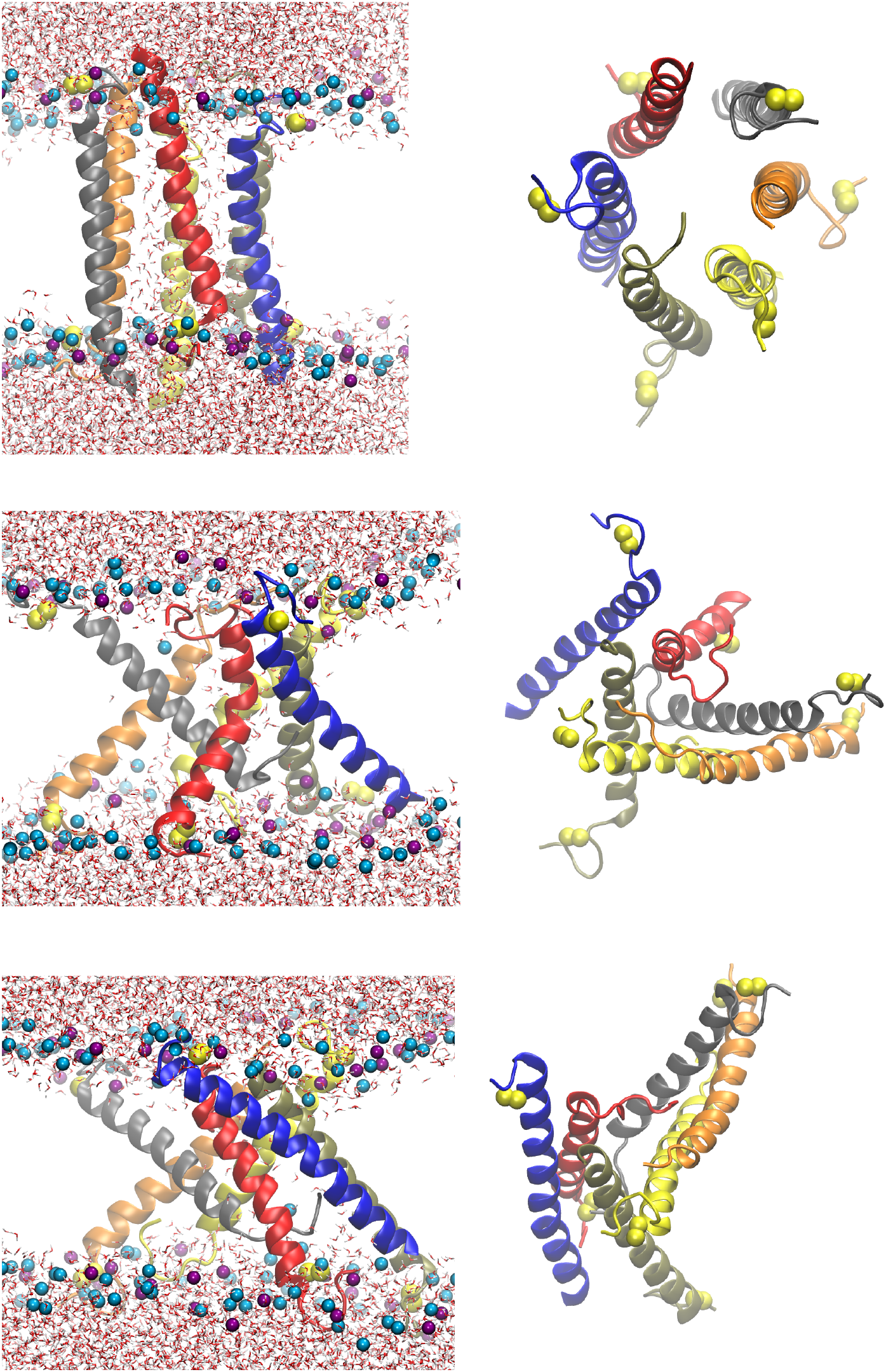
Antiparallel IAPP helices in initial configuration (above), after 2 μs of the simulation (middle), and at the end of the simulation (10 μs, lower). Left panels side views, right panels top views. Color code as in Fig. 3.

The configuration of the parallel β-barrel is highly stable over 10 μs of the simulation with movements of the termini highly restricted (Figs. 5a and 5b). On the other hand, the antiparallel β-barrel collapsed after 1 μs and consequently the initial pore dissipated (Figs. 6a and 6b). So, this simulation was not continued further.

**Fig. 5.**
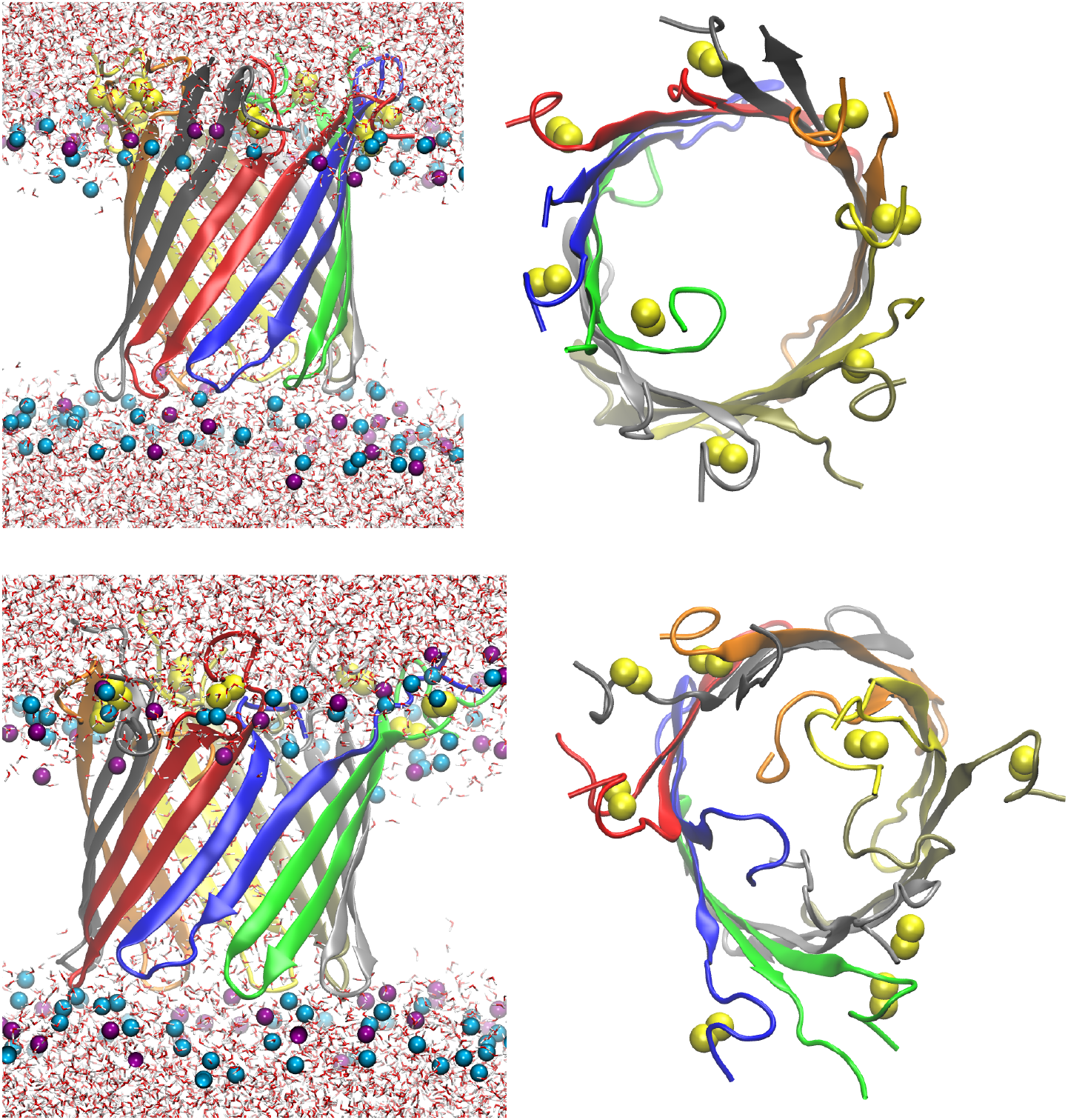
IAPP parallel β-barrel in initial configuration (top) and at the end of the simulation (bottom). Side views (left) and top views (right). Color code same as in Fig. 3.

**Fig. 6.**
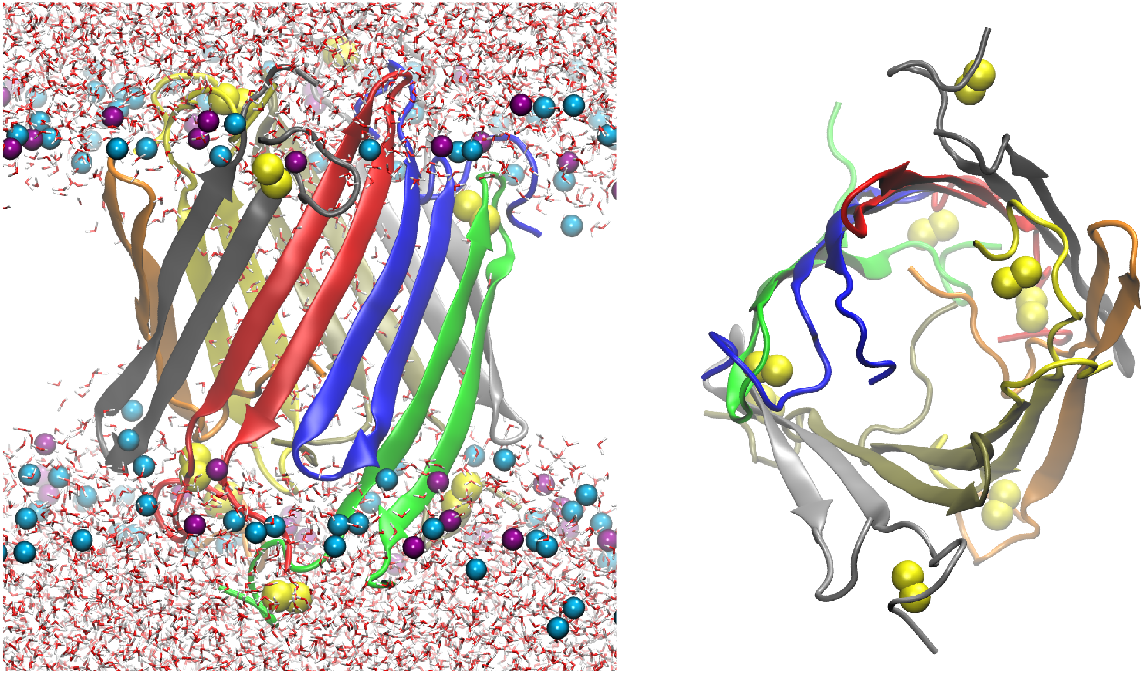
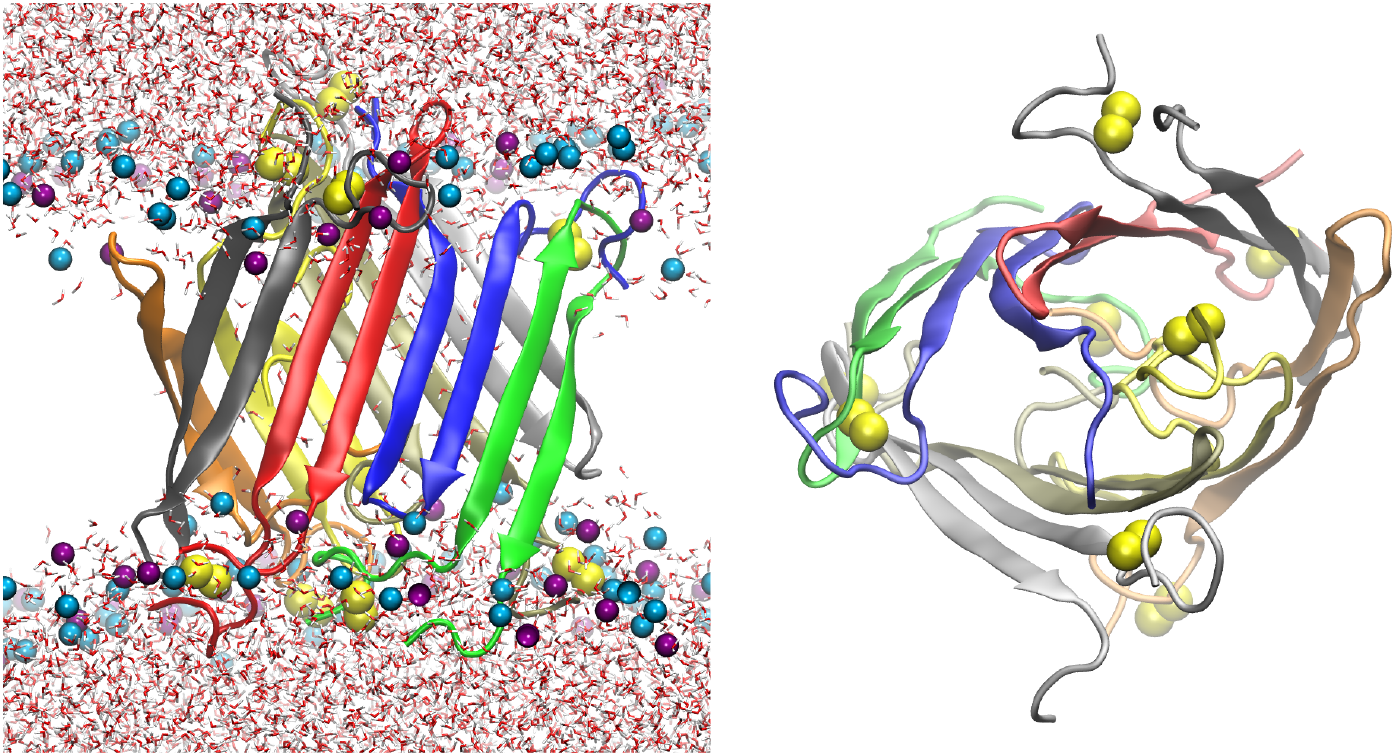
IAPP antiparallel β-barrel in initial configuration (top) and after 1 us of simulation (bottom). Side views (left) and top views (right). Color code same as in Fig. 3.

The pore radius results are shown in Fig. 7. The parallel helical configuration gives a slightly larger pore than for the antiparallel one. However, the pore size seems to be more stable in antiparallel configuration, while it fluctuates highly in the parallel one. Compared to a potent pore-forming peptide, such as macrolittin70 ^67^, IAPP long helices failed to produce a large pore. This is probably due to the lack of proper interactions to keep adjacent peptides tilted towards each other. The pore size for the octameric parallel β-barrel is large and very stable throughout the simulation.

**Fig. 7.**
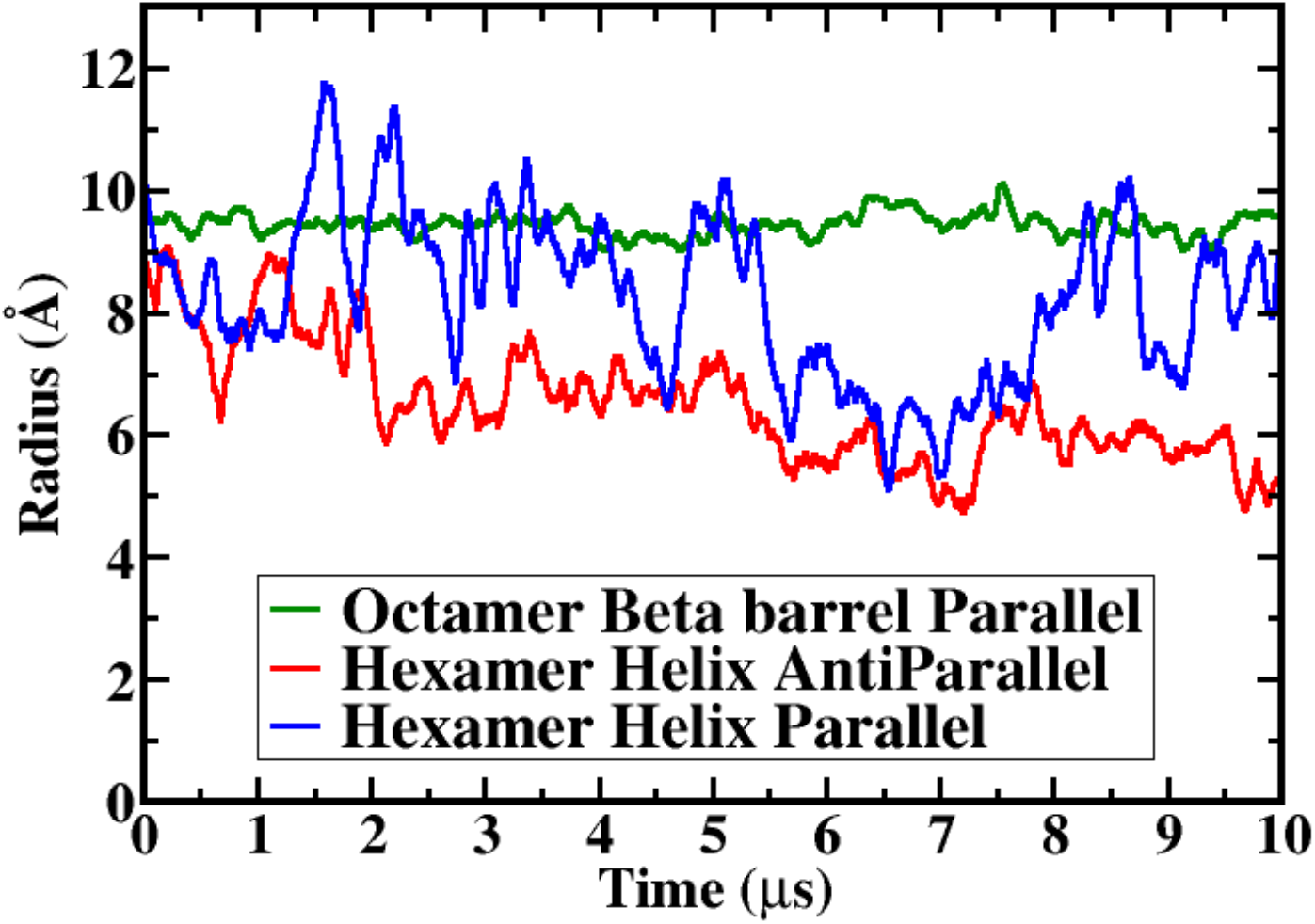
Pore radius as a function of simulation time.

There are 15 peptide pairs in a hexamer and 28 pairs in an octamer. Table 3 lists the 10 strongest peptide pair interaction energies in each system calculated over the last 2 μs of each simulation. For a potent peptide, it is expected that pair interactions between adjacent peptides are stronger than non-adjacent ones. This is true for the parallel β-barrel but not true for the helices.

**Table 3.** 10 strongest protein-protein interaction energies (kcal/mol) for different systems of IAPP calculated over the last 2 μs of each simulation.

| | Parallel Helices | Antiparallel Helices | Parallel β-barrel |
| --- | --- | --- | --- |
| 1 | D-E | D-E | C-D |
| | -79.2 ± 6.6 | -124.2 ± 4.1 | -154.6 ± 3.9 |
| 2 | B-C | A-B | G-H |
| | -55.5 ± 6.2 | -93.7 ± 3.5 | -142.3 ± 3.4 |
| 3 | C-D | B-C | E-F |
| | -54.0 ± 3.6 | -91.5 ± 3.7 | -136.7 ± 2.9 |
| 4 | B-D | E-F | A-B |
| | -22.5 ± 2.8 | -33.6 ± 2.2 | -131.3 ± 7.8 |
| 5 | C-E | B-D | D-E |
| | -21.4 ± 3.7 | -25.0 ± 0.6 | -130.5 ± 2.9 |
| 6 | A-F | C-E | A-H |
| | -19.3 ± 0.9 | -24.5 ± 0.3 | -114.3 ± 8.4 |
| 7 | A-B | C-D | F-G |
| | -13.9 ± 2.0 | -24.4 ± 0.3 | -110.6 ± 6.0 |
| 8 | A-C | A-F | B-C |
| | -13.1 ± 4.5 | -23.8 ± 0.6 | -106.9 ± 4.1 |
| 9 | E-F | B-F | A-G |
| | -12.7 ± 2.1 | -22.0 ± 0.9 | -18.8 ± 5.4 |
| 10 | A-E | C-F | F-H |
| | -12.1 ± 3.4 | -18.5 ± 2.5 | -7.5 ± 2.6 |

Two observations are possible in the interaction energies for helices. First, strong non-adjacent interactions occur between peptides with at least one unfolded C-terminus e.g., B-D and C-E. Second, there are stronger adjacent interactions in the antiparallel configuration than in the parallel one. This is probably due to favorable interactions between unlike termini. The fluctuations in pore size of parallel configuration is possibly because of repulsive interactions between non-adjacent C-termini. The second observation is also prominent in the parallel β-barrel. In this configuration, the C-terminus of each peptide is close to the N-terminus of the adjacent peptide which causes favorable interactions that likely stabilize the pore. Table 4 shows that the strongest residue-residue interactions in the parallel β-barrel are between N22-H18, T30-Q10, R11-S28 or S29, and N31-T9.

**Table 4.** The three strongest residue-residue interactions in each adjacent molecular pair in the parallel β-barrel calculated over the last 2 μs of the simulation (kcal/mol).

| A-B | B-C | C-D | D-E | E-F | F-G | G-H | H-A |
| --- | --- | --- | --- | --- | --- | --- | --- |
| AN22-BH18 | BT30-CQ10 | CN22-DH18 | DS29-ER11 | ET30-FQ10 | FS29-GR11 | GS29-HR11 | HN22-AH18 |
| -7.18 ± 0.05 | -5.83 ± 0.31 | -7.59 ± 0.32 | -8.11 ± 0.56 | -6.61 ± 0.15 | -6.67 ± 0.67 | -10.89 ± 0.55 | -8.30 ± 0.14 |
| AT30-BQ10 | BN22-CH18 | CT30-DQ10 | DT30-EQ10 | ES29-FR11 | FN22-GH18 | GN22-HH18 | HT30-AQ10 |
| -6.88 ± 0.08 | -4.77 ± 0.19 | -7.09 ± 0.27 | -6.30 ± 0.07 | -5.76 ± 1.46 | -5.48 ± 0.23 | -7.34 ± 0.57 | -5.22 ± 1.32 |
| AS29-BR11 | BS28-CR11 | CS28-DR11 | DS28-ER11 | EN31-FT9 | FS28-GR11 | GT30-HQ10 | HS28-AR11 |
| -5.52 ± 1.87 | -4.50 ± 0.20 | -4.88 ± 0.06 | -5.79 ± 0.21 | -5.55 ± 0.49 | -4.86 ± 0.10 | -6.82 ± 0.09 | -4.49 ± 0.48 |

Protein-lipid interaction energies are listed in Table 5. We have calculated these data over the first 2 μs and the last 2 μs of each simulation to see their changes with time. Note that the POPC:POPG ratio is 7:3, so, there are more POPC lipids to interact with peptides. Also note that during the first 2 μs of simulations of IAPP helices, peptides tilt or unfold their C-terminus. So, our results are affected both by this and lipid arrangements. In all cases, protein-POPC interactions become weaker which shows larger distances between peptides and POPC lipids. For helical configurations, protein-POPG interactions do not show significant changes over time and their differences are within statistical error. However, in β-barrel parallel configuration, protein-POPG interactions become stronger especially on the upper leaflet where the termini reside. This shows more packed arrangement of POPG around peptides at the end of the simulation.

**Table 5.** Protein-lipid interaction energies (kcal/mol) for different systems of IAPP.

| Configuration | Leaflet | Protein-POPC |  | Protein-POPG |  |
| --- | --- | --- | --- | --- | --- |
| | | First 2 μs | Last 2 μs | First 2 μs | Last 2 μs |
| Helix Parallel | Lower leaflet | -329 ± 26 | -252 ± 10 | -234 ± 12 | -226 ± 24 |
| | Upper leaflet | -820 ± 33 | -689 ± 19 | -772 ± 24 | -755 ± 35 |
| Helix Antiparallel | Lower leaflet | -580 ± 44 | -544 ± 20 | -548 ± 38 | -501 ± 14 |
| | Upper leaflet | -715 ± 58 | -410 ± 22 | -404 ± 20 | -499 ± 27 |
| β-barrel Parallel | Lower leaflet | -371 ± 22 | -322 ± 22 | -139 ± 23 | -178 ± 20 |
| | Upper leaflet | -746 ± 16 | -703 ± 12 | -740 ± 19 | -936 ± 49 |

## DISCUSSION

This work evaluates possible membrane pore structures formed by oligomers of the IAPP peptide implicated in type II diabetes. We find that transmembrane helical bundles are stably embedded in a mixed POPC/POPG membrane on a 10-μs timescale but make narrow water pores. Partial unfolding at the C-terminus and/or tilting of the helices induce interactions between nonadjacent peptides that block the channel. A β-barrel consisting of parallel β-hairpins at residues 11-30 is very stable, whereas the corresponding barrel with antiparallel hairpins is not. The number of monomers in these structures is arbitrary; six helices were chosen for comparison with previous work on AMPs and eight monomers were chosen for the barrels to provide a large enough channel and avoid steric clashes in the interior. It is expected that slightly smaller or larger oligomers will exhibit similar stability.

We note that IAPP binds an aggregation inhibitor as a β-hairpin similar but not identical to that in our β-barrel models ^77^. There is more twist in that structure and a different h-bonding pattern. Similar hairpins have been frequently observed in MD simulations of the monomer ^78–80^, showing that such conformations are easily accessible and thermodynamically stable. The high kinetic stability of the parallel β-barrel does not prove that this structure occurs in reality. It could correspond to a deep local free energy minimum that may not be competitive with other minima and/or kinetically accessible.

The picture that emerges from our results together with published experimental data is that different pore structures may occur under different conditions. A monomeric helical peptide can act like helical AMPs but an aggregated β-sheet oligomer may disrupt membranes by inserting into them an oligomer of β hairpins. The lack of perfect correlation between liposome leakage and cell toxicity^24^ may be due to the fact that the former is dominated by helical bundles whereas the latter by β sheet oligomers. Barrels may be more physiologically relevant because the requirement of anionic membrane is less severe and pancreatic cell membranes are not highly anionic^81^. The finding that off-register β sheet oligomers are more toxic than in-register ones^82^ is consistent with the β-barrel model, because β-barrels require shear for stability^66^, and shear is equivalent to off-register β sheets.

Most computational work on IAPP has focused on aggregation in solution ^47,48^. A smaller number of articles considered interactions with membranes ^19,49–51,54,55,83^. Zhao et al. ^84,85^ assumed that the peptide conformation is the same as in the fibril ^29^, similar to previous work on amyloid β ^86^. In these pore structures the peptides adopt a U-shaped conformation and make a double barrel with two concentric cylindrical β-sheets. Such double barrels have never been experimentally observed. The longer distance between strands in the outer barrel makes it very difficult for them to maintain h bonding; as a result the barrel breaks into smaller oligomers. Such models are not stable in our implicit membrane pores (results not shown). In addition, fibril structures vary considerably ^32–34^ and there is evidence that the peptide conformation is different between fibrils and toxic oligomers ^11,14^. More recent models of Aβ pores are based on classical, single-layer barrels, for the full-length ^87^ or the 25-35 fragment ^88^. These are similar in concept to the present barrels for IAPP. The difference in the present work is that we pre-evaluate the models using an implicit membrane model, which, in our experience, can be more discriminating than all-atom simulations. A couple of coarse-grained simulation studies treated the peptide as helical and obtained pore-like aggregates under certain conditions ^52,53^. These are similar to our helical bundles but the present models are of course more atomically detailed and allow changes in secondary structure.

Since the pore models are hypothetical, it is imperative to examine them in the light of available experimental data. Ramamoorthy and coworkers found that the 1-19 fragment of IAPP is helical and causes dye leakage from POPG liposomes without forming amyloid fibrils^46^. This finding seems consistent with our helical bundles, in which there is a tendency for unfolding at the Ct, while the segment 9-20 tends to remain helical and appears capable of supporting a pore at low tilt angle (Figs. 3,4). The experimental observation that the full-length peptide is less effective in inducing dye leakage than the 1-19 fragment under these conditions could be explained by the blocking of the channel by the unfolded C-termini we observe in the simulations. The partially unfolded helices in our helical bundles are similar to the structure of IAPP in micelles^57,58^. The helical bundles are also consistent with the experiments of Miranker and coworkers^16,89^. Antiparallel dimers like those seen in Fig. 4 were observed by FRET on DOPG nanodiscs^90^. Some electrophysiology studies gave uniform single-channel conductances consistent with a well-defined pore structure ^39,42^ but others did not^9,41,91^]. A SFG study of IAPP gave a tilt angle of the β strands similar to what is observed in our β barrel model^92^.

The rat analogue of IAPP (rIAPP) is not toxic ^93,94^, although some *in vitro* studies found some membrane disruption at higher concentrations ^24,95,96^. rIAPP differs from hIAPP at 6 sites between 18 and 29 (H18R,F23L,A25P,I26V,S28P,S29P). This would place at least two prolines in a β strand, which would be clearly unfavorable. In addition, Arg in the interior of a barrel may be less stabilizing than His due to steric and electrostatic repulsion. So, the rat mutations destabilize not only the fibril but also a β barrel putative membrane pore structure. Analogues from other species that are nontoxic (bovine)^97^ or less toxic (ursine)^98^ also tend to have at least one or two P in the 11-30 region. These mutations should be more tolerable in the helical bundles, which may explain the observation of significant liposome leakage activity of rIAPP in some experiments ^16,89,99^. The helix bundles have similarities to those of magainin ^60^, corroborating the experimental comparison between the two peptides ^16^.

Validating these structures experimentally is challenging, but progress is being made on several fronts. Direct structure determination of the barrel state might be possible if heterogeneity was suppressed. To do that, one could synthesize a tandem octamer of IAPP and try to fold and crystallize in various membrane mimicking environments. Indirect tests, such as mutations, are also possible. For example, the turns in the β barrel structure should tolerate mutation to P. To our knowledge, the S20P mutant has been considered only in the context of the 11-25 fragment and was found to not self-assemble^100^. N21P was found to accelerate self-assembly and to be as toxic as wild type, but N21G was not toxic^101^. Other possible approaches include cryoET, which has been recently applied to Ab oligomers^102^. Interestingly, when de novo designed β barrels were too stable they were found to form amyloid^76^, showing a link between these two states. Even if these β barrels do not actually occur in Nature, it would be important to understand why, given their high stability *in silico*.

The only early-onset diabetes II mutation in IAPP is S20G, observed in Asian populations ^103^. It is known to accelerate fibril formation^104,105^ and to be more toxic^106^. In the barrel structure S20 is on the turn. It does not affect much the stability of the barrel but may facilitate the formation of the hairpin. Another possibility is that, given that the fibril structure of the mutant is different ^33^ it may experience a lower barrier to dissociation from the fibril and thus be more available for interactions with membranes. In the helical conformation S20 (as well as S19 and T30) are on the hydrophobic face of the helix and mutation to G likely increases the affinity for lipids and thus the stability of the pore. An interesting question is whether S19G would have similar effects as S20G.

Fragments of many amyloidogenic peptides have been found to be toxic. Such is the case for fragment 20-29 of IAPP^6^. This fragment forms one strand of the hairpin studied here and could presumably form a barrel by itself. However, shorter fragments that were found to be toxic in the fibril form, but not freshly dissolved, such as 23-27, are harder to rationalize in this way and need further investigation. Models of such fragments will be constructed and evaluated in the near future. The ΦχΦχΦ pattern required for membrane-embedded β barrels should be widespread in protein sequence space, which may explain the observed toxicity of prefibrillar aggregates of a variety of proteins^107^.

## Acknowledgments

This work was supported by the NSF (MCB 1855942). Anton computer time was provided by the Pittsburgh Supercomputing Center through grant R01GM116961 from the NIH. The Anton machine at PSC was generously made available by D.E. Shaw Research, and computer time was provided by the National Center for Multiscale Modeling of Biological Systems through grant number P41GM103712-S1 from the NIH and Pittsburgh Supercomputing Center.

## Author Contributions

TL, AS and BN performed the simulations and wrote the manuscript.

## Notes

### Competing Interest Statement

The authors have declared no competing interest.

## References

(1) Mukherjee, A.; Morales-Scheihing, D.; Butler, P. C.; Soto, C. Type 2 Diabetes as a Protein Misfolding Disease. Trends Mol. Med. 2015, 21, 439–449.

(2) Hebda, J. A.; Miranker, A. D. The Interplay of Catalysis and Toxicity by Amyloid Intermediates on Lipid Bilayers: Insights from Type II Diabetes. Annu. Rev. Biophys. 2009, 38, 125–152.

(3) Cao, P.; Abedini, A.; Raleigh, D. P. Aggregation of Islet Amyloid Polypeptide: From Physical Chemistry to Cell Biology. Curr. Opin. Struct. Biol. 2013, 23, 82–89.

(4) Patel, H. R.; Pithadia, A. S.; Brender, J. R.; Fierke, C. A.; Ramamoorthy, A. In Search of Aggregation Pathways of IAPP and Other Amyloidogenic Proteins: Finding Answers through NMR Spectroscopy. J. Phys. Chem. Lett. 2014, 5, 1864–1870.

(5) Lorenzo, A.; Razzaboni, B.; Weir, G. C.; Yankner, B. A. Pancreatic Islet Cell Toxicity of Amylin Associated with Type-2 Diabetes Mellitus. Nature 1994, 368, 756–760.

(6) Tenidis, K.; Waldner, M.; Bernhagen, J.; Fischle, W.; Bergmann, M.; Weber, M.; Merkle, M. L.; Voelter, W.; Brunner, H.; Kapurniotu, A. Identification of a Penta- and Hexapeptide of Islet Amyloid Polypeptide (IAPP) with Amyloidogenic and Cytotoxic Properties. J. Mol. Biol. 2000, 295, 1055–1071.

(7) Krotee, P.; Rodriguez, J. A.; Sawaya, M. R.; Cascio, D.; Reyes, F. E.; Shi, D.; Hattne, J.; Nannenga, B. L.; Oskarsson, M. E.; Philipp, S.; et al. Atomic Structures of Fibrillar Segments of HIAPP Suggest Tightly Mated β-Sheets Are Important for Cytotoxicity. Elife 2017, 6, 1–26.

(8) Pilkington, E. H.; Gurzov, E. N.; Kakinen, A.; Litwak, S. A.; Stanley, W. J.; Davis, T. P.; Ke, P. C. Pancreatic β-Cell Membrane Fluidity and Toxicity Induced by Human Islet Amyloid Polypeptide Species. Sci. Rep. 2016, 6, 1–10.

(9) Janson, J.; Ashley, R. H.; Harrison, D.; McIntyre, S.; Butler, P. C. The Mechanism of Islet Amyloid Polypeptide Toxicity Is Membrane Disruption by Intermediate-Sized Toxic Amyloid Particles. Diabetes 1999, 48, 491–498.

(10) Anguiano, M.; Nowak, R. J.; Lansbury, P. T. Protofibrillar Islet Amyloid Polypeptide Permeabilizes Synthetic Vesicles by a Pore-like Mechanism That May Be Relevant to Type II Diabetes. Biochemistry 2002, 41, 11338–11343.

(11) Konarkowska, B.; Aitken, J. F.; Kistler, J.; Zhang, S.; Cooper, G. J. S. The Aggregation Potential of Human Amylin Determines Its Cytotoxicity towards Islet β-Cells. FEBS J. 2006, 273, 3614–3624.

(12) Weise, K.; Radovan, D.; Gohlke, A.; Opitz, N.; Winter, R. Interaction of HIAPP with Model Raft Membranes and Pancreatic β-Cells: Cytotoxicity of HIAPP Oligomers. ChemBioChem 2010, 11, 1280–1290.

(13) Ridgway, Z.; Lee, K. H.; Zhyvoloup, A.; Wong, A.; Eldrid, C.; Hannaberry, E.; Thalassinos, K.; Abedini, A.; Raleigh, D. P. Analysis of Baboon IAPP Provides Insight into Amyloidogenicity and Cytotoxicity of Human IAPP. Biophys. J. 2020, 118, 1142–1151.

(14) Abedini, A.; Plesner, A.; Cao, P.; Ridgway, Z.; Zhang, J.; Tu, L. H.; Middleton, C. T.; Chao, B.; Sartori, D. J.; Meng, F.; et al. Time-Resolved Studies Define the Nature of Toxic IAPP Intermediates, Providing Insight for Anti-Amyloidosis Therapeutics. Elife 2016, 5, 1–28.

(15) Jayasinghe, S. A.; Langen, R. Membrane Interaction of Islet Amyloid Polypeptide. Biochim. Biophys. Acta - Biomembr. 2007, 1768, 2002–2009.

(16) Last, N. B.; Miranker, A. D. Common Mechanism Unites Membrane Poration by Amyloid and Antimicrobial Peptides. Proc Natl Acad Sci U S A 2013, 110, 6382–6387.

(17) Zhang, X.; St Clair, J. R.; London, E.; Raleigh, D. P. Islet Amyloid Polypeptide Membrane Interactions: Effects of Membrane Composition. Biochemistry 2017, 56, 376–390.

(18) Birol, M.; Kumar, S.; Rhoades, E.; Miranker, A. D. Conformational Switching within Dynamic Oligomers Underpins Toxic Gain-of-Function by Diabetes-Associated Amyloid. Nat. Commun. 2018, 9, 1–12.

(19) Martel, A.; Antony, L.; Gerelli, Y.; Porcar, L.; Fluitt, A.; Hoffmann, K.; Kiesel, I.; Vivaudou, M.; Fragneto, G.; De Pablo, J. J. Membrane Permeation versus Amyloidogenicity: A Multitechnique Study of Islet Amyloid Polypeptide Interaction with Model Membranes. J. Am. Chem. Soc. 2017, 139, 137–148.

(20) Sparr, E.; Engel, M. F. M.; Sakharov, D. V.; Sprong, M.; Jacobs, J.; De Kruijff, B.; Höppener, J. W. M.; Antoinette Killian, J. Islet Amyloid Polypeptide-Induced Membrane Leakage Involves Uptake of Lipids by Forming Amyloid Fibers. FEBS Lett. 2004, 577, 117–120.

(21) Domanov, Y. A.; Kinnunen, P. K. J. Islet Amyloid Polypeptide Forms Rigid Lipid-Protein Amyloid Fibrils on Supported Phospholipid Bilayers. J. Mol. Biol. 2008, 376, 42–54.

(22) Engel, M. F. M.; Khemtémourian, L.; Kleijer, C. C.; Meeldijk, H. J. D.; Jacobs, J.; Verkleij, A. J.; De Kruijff, B.; Killian, J. A.; Höppener, J. W. M. Membrane Damage by Human Islet Amyloid Polypeptide through Fibril Growth at the Membrane. Proc. Natl. Acad. Sci. U. S. A. 2008, 105, 6033–6038.

(23) Haataja, L.; Gurlo, T.; Huang, C. J.; Butler, P. C. Islet Amyloid in Type 2 Diabetes, and the Toxic Oligomer Hypothesis. Endocr. Rev. 2008, 29, 303–316.

(24) Cao, P.; Abedini, A.; Wang, H.; Tu, L. H.; Zhang, X.; Schmidt, A. M.; Raleigh, D. P. Islet Amyloid Polypeptide Toxicity and Membrane Interactions. Proc. Natl. Acad. Sci. U. S. A. 2013, 110, 19279–19284.

(25) Hamley, I. W. The Amyloid Beta Peptide: A Chemist’s Perspective. Role in Alzheimer’s and Fibrillization. Chem. Rev. 2012, 112, 5147–5192.

(26) Meade, R. M.; Fairlie, D. P.; Mason, J. M. Alpha-Synuclein Structure and Parkinson’s Disease - Lessons and Emerging Principles. Mol. Neurodegener. 2019, 14, 1–14.

(27) Jayasinghe, S. A.; Langen, R. Identifying Structural Features of Fibrillar Islet Amyloid Polypeptide Using Site-Directed Spin Labeling. J. Biol. Chem. 2004, 279, 48420–48425.

(28) Wiltzius, J. J. W.; Sievers, S. A.; Sawaya, M. R.; Cascio, D.; Popov, D.; Riekel, C.; Eisenberg, D. Atomic Structure of the Cross-β Spine of Islet Amyloid Polypeptide (Amylin). Protein Sci. 2008, 17, 1467–1474.

(29) Luca, S.; Yau, W. M.; Leapman, R.; Tycko, R. Peptide Conformation and Supramolecular Organization in Amylin Fibrils: Constraints from Solid-State NMR. Biochemistry 2007, 46, 13505–13522.

(30) Kajava, A. V.; Aebi, U.; Steven, A. C. The Parallel Superpleated Beta-Structure as a Model for Amyloid Fibrils of Human Amylin. J. Mol. Biol. 2005, 348, 247–252.

(31) Röder, C.; Kupreichyk, T.; Gremer, L.; Schäfer, L.; Pothula, K.; Ravelli, R.; Willbold, D.; Hoyer, W.; Schröder, G. Amyloid Fibril Structure of Islet Amyloid Polypeptide by Cryo-Electron Microscopy Reveals Similarities with Amyloid Beta. 2020.

(32) Cao, Q.; Boyer, D. R.; Sawaya, M. R.; Ge, P.; Eisenberg, D. S. Cryo-EM Structure and Inhibitor Design of Human IAPP (Amylin) Fibrils. Nat. Struct. Mol. Biol. 2020, 27, 653–659.

(33) Gallardo, R.; Iadanza, M. G.; Xu, Y.; Heath, G. R.; Foster, R.; Radford, S. E.; Ranson, N. A. Fibril Structures of Diabetes-Related Amylin Variants Reveal a Basis for Surface-Templated Assembly. Nat. Struct. Mol. Biol. 2020, 27, 1048–1056.

(34) Röder, C.; Kupreichyk, T.; Gremer, L.; Schäfer, L. U.; Pothula, K. R.; Ravelli, R. B. G.; Willbold, D.; Hoyer, W.; Schröder, G. F. Cryo-EM Structure of Islet Amyloid Polypeptide Fibrils Reveals Similarities with Amyloid-β Fibrils. Nat. Struct. Mol. Biol. 2020, 27, 660–667.

(35) Soriaga, A. B.; Sangwan, S.; MacDonald, R.; Sawaya, M. R.; Eisenberg, D. Crystal Structures of IAPP Amyloidogenic Segments Reveal a Novel Packing Motif of Out-of-Register Beta Sheets. J. Phys. Chem. B 2016, 120, 5810–5816.

(36) Laganowsky, A.; Liu, C.; Sawaya, M. R.; Whitelegge, J. P.; Park, J.; Zhao, M.; Pensalfini, A.; Soriaga, A. B.; Landau, M.; Teng, P. K.; et al. Atomic View of a Toxic Amyloid Small Oligomer. Science 2012, 335, 1228–1231.

(37) Rawat, A.; Maity, B. K.; Chandra, B.; Maiti, S. Aggregation-Induced Conformation Changes Dictate Islet Amyloid Polypeptide (IAPP) Membrane Affinity. Biochim. Biophys. Acta - Biomembr. 2018, 1860, 1734–1740.

(38) Rodriguez Camargo, D. C.; Korshavn, K. J.; Jussupow, A.; Raltchev, K.; Goricanec, D.; Fleisch, M.; Sarkar, R.; Xue, K.; Aichler, M.; Mettenleiter, G.; et al. Stabilization and Structural Analysis of a Membrane-Associated HIAPP Aggregation Intermediate. Elife 2017, 6, 1–22.

(39) Mirzabekov, T. A.; Lin, M. C.; Kagan, B. L. Pore Formation by the Cytotoxic Islet Amyloid Peptide Amylin. J. Biol. Chem. 1996, 271, 1988–1992.

(40) Porat, Y.; Kolusheva, S.; Jelinek, R.; Gazit, E. The Human Islet Amyloid Polypeptide Forms Transient Membrane-Active Prefibrillar Assemblies. Biochemistry 2003, 42, 10971–10977.

(41) Kayed, R.; Sokolov, Y.; Edmonds, B.; McIntire, T. M.; Milton, S. C.; Hall, J. E.; Glabe, C. G. Permeabilization of Lipid Bilayers Is a Common Conformation-Dependent Activity of Soluble Amyloid Oligomers in Protein Misfolding Diseases. J. Biol. Chem. 2004, 279, 46363–46366.

(42) Quist, A.; Doudevski, I.; Lin, H.; Azimova, R.; Ng, D.; Frangione, B.; Kagan, B.; Ghiso, J.; Lal, R. Amyloid Ion Channels: A Common Structural Link for Protein-Misfolding Disease. Proc Natl Acad Sci U S A 2005, 102, 10427–10432.

(43) Demuro, A.; Mina, E.; Kayed, R.; Milton, S. C.; Parker, I.; Glabe, C. G. Calcium Dysregulation and Membrane Disruption as a Ubiquitous Neurotoxic Mechanism of Soluble Amyloid Oligomers. J. Biol. Chem. 2005, 280, 17294–17300.

(44) Lu, T.; Meng, F.; Wei, Y.; Li, Y.; Wang, C.; Li, F. Exploring the Relation between the Oligomeric Structure and Membrane Damage by a Study on Rat Islet Amyloid Polypeptide. Phys. Chem. Chem. Phys. 2018, 20, 8976–8983.

(45) Green, J. D.; Kreplak, L.; Goldsbury, C.; Li Blatter, X.; Stolz, M.; Cooper, G. S.; Seelig, A.; Kistler, J.; Aebi, U. Atomic Force Microscopy Reveals Defects within Mica Supported Lipid Bilayers Induced by the Amyloidogenic Human Amylin Peptide. J. Mol. Biol. 2004, 342, 877–887.

(46) Brender, J. R.; Lee, E. L.; Cavitt, M. A.; Gafni, A.; Steel, D. G.; Ramamoorthy, A. Amyloid Fiber Formation and Membrane Disruption Are Separate Processes Localized in Two Distinct Regions of IAPP, the Type-2-Diabetes-Related Peptide. J. Am. Chem. Soc. 2008, 130, 6424–6429.

(47) Morriss-Andrews, A.; Shea, J. E. Simulations of Protein Aggregation: Insights from Atomistic and Coarse-Grained Models. J. Phys. Chem. Lett. 2014, 5, 1899–1908.

(48) Dong, X.; Qiao, Q.; Qian, Z.; Wei, G. Recent Computational Studies of Membrane Interaction and Disruption of Human Islet Amyloid Polypeptide: Monomers, Oligomers and Protofibrils. Biochim. Biophys. Acta - Biomembr. 2018, 1860, 1826–1839.

(49) Duan, M.; Fan, J.; Huo, S. Conformations of Islet Amyloid Polypeptide Monomers in a Membrane Environment: Implications for Fibril Formation. PLoS One 2012, 7.

(50) Zhang, Y.; Luo, Y.; Deng, Y.; Mu, Y.; Wei, G. Lipid Interaction and Membrane Perturbation of Human Islet Amyloid Polypeptide Monomer and Dimer by Molecular Dynamics Simulations. PLoS One 2012, 7.

(51) Dignon, G. L.; Zerze, G. H.; Mittal, J. Interplay between Membrane Composition and Structural Stability of Membrane-Bound HIAPP. J. Phys. Chem. B 2017, 121, 8661–8668.

(52) Xu, W.; Wei, G.; Su, H.; Nordenskiöld, L.; Mu, Y. Effects of Cholesterol on Pore Formation in Lipid Bilayers Induced by Human Islet Amyloid Polypeptide Fragments: A Coarse-Grained Molecular Dynamics Study. Phys. Rev. E - Stat. Nonlinear, Soft Matter Phys. 2011, 84, 1–8.

(53) Pannuzzo, M.; Raudino, A.; Milardi, D.; La Rosa, C.; Karttunen, M. α-Helical Structures Drive Early Stages of Self-Assembly of Amyloidogenic Amyloid Polypeptide Aggregate Formation in Membranes. Sci. Rep. 2013, 3, 1–10.

(54) Poojari, C.; Xiao, D.; Batista, V. S.; Strodel, B. Membrane Permeation Induced by Aggregates of Human Islet Amyloid Polypeptides. Biophys. J. 2013, 105, 2323–2332.

(55) Qian, Z.; Zou, Y.; Zhang, Q.; Chen, P.; Ma, B.; Wei, G.; Nussinov, R. Atomistic-Level Study of the Interactions between HIAPP Protofibrils and Membranes: Influence of PH and Lipid Composition. Biochim. Biophys. Acta - Biomembr. 2018, 1860, 1818–1825.

(56) Apostolidou, M.; Jayasinghe, S. A.; Langen, R. Structure of α-Helical Membrane-Bound Human Islet Amyloid Polypeptide and Its Implications for Membrane-Mediated Misfolding. J. Biol. Chem. 2008, 283, 17205–17210.

(57) Patil, S. M.; Xu, S.; Sheftic, S. R.; Alexandrescu, A. T. Dynamic α-Helix Structure of Micelle-Bound Human Amylin. J. Biol. Chem. 2009, 284, 11982–11991.

(58) Nanga, R. P. R.; Brender, J. R.; Xu, J.; Hartman, K.; Subramanian, V.; Ramamoorthy, A. Three-Dimensional Structure and Orientation of Rat Islet Amyloid Polypeptide Protein in a Membrane Environment by Solution NMR Spectroscopy. J. Am. Chem. Soc. 2009, 131, 8252–8261.

(59) Soong, R.; Brender, J. R.; Macdonald, P. M.; Ramamoorthy, A. Association of Highly Compact Type Il Diabetes Related Islet Amyloid Polypeptide Intermediate Species at Physiological Temperature Revealed by Diffusion NMR Spectroscopy. J. Am. Chem. Soc. 2009, 131, 7079–7085.

(60) Pino-Angeles, A.; Leveritt, J. M. I.; Lazaridis, T. Pore Structure and Synergy in Antimicrobial Peptides of the Magainin Family. PLoS Comput Biol 2016, 12, e1004570.

(61) Lazaridis, T. Effective Energy Function for Proteins in Lipid Membranes. Proteins Struct. Funct. Bioinforma. 2003, 52, 176–192.

(62) Lazaridis, T.; Karplus, M. Effective Energy Function for Proteins in Solution. Proteins Struct. Funct. Bioinforma. 1999, 35, 133–152.

(63) Lazaridis, T. Implicit Solvent Simulations of Peptide Interactions with Anionic Lipid Membranes. Proteins-Structure Funct. Bioinforma. 2005, 58, 518–527.

(64) Mihajlovic, M.; Lazaridis, T. Antimicrobial Peptides Bind More Strongly to Membrane Pores. Biochim. Biophys. Acta-Biomembranes 2010, 1798, 1494–1502.

(65) Lazaridis, T. Structural Determinants of Transmembrane Beta-Barrels. J. Chem. Theory Comput. 2005, 1, 716–722.

(66) Lipkin, R.; Pino-Angeles, A.; Lazaridis, T. Transmembrane Pore Structures of β-Hairpin Antimicrobial Peptides by All-Atom Simulations. J. Phys. Chem. B 2017, 121, 9126–9140.

(67) Sepehri, A.; PeBenito, L.; Pino-Angeles, A.; Lazaridis, T. What Makes a Good Pore Former: A Study of Synthetic Melittin Derivatives. Biophys. J. 2020, 118, 1901–1913.

(68) Jo, S.; Lim, J. B.; Klauda, J. B.; Im, W. CHARMM-GUI Membrane Builder for Mixed Bilayers and Its Application to Yeast Membranes. Biophys. J. 2009, 97, 50–58.

(69) Phillips, J. C.; Braun, R.; Wang, W.; Gumbart, J.; Tajkhorshid, E.; Villa, E.; Chipot, C.; Skeel, R. D.; Kale, L.; Schulten, K. Scalable Molecular Dynamics with NAMD. J. Comput. Chem. 2005, 26, 1781–1802.

(70) Mihajlovic, M.; Lazaridis, T. Membrane-Bound Structure and Energetics of Alpha-Synuclein. Proteins-Structure Funct. Bioinforma. 2008, 70, 761–778.

(71) McLean, L. R.; Balasubramaniam, A. Promotion of β-Structure by Interaction of Diabetes Associated Polypeptide (Amylin) with Phosphatidylcholine. Biochim. Biophys. Acta (BBA)/Protein Struct. Mol. 1992, 1122, 317–320.

(72) Knight, J. D.; Miranker, A. D. Phospholipid Catalysis of Diabetic Amyloid Assembly. J. Mol. Biol. 2004, 341, 1175–1187.

(73) Jayasinghe, S. A.; Langen, R. Lipid Membranes Modulate the Structure of Islet Amyloid Polypeptide. Biochemistry 2005, 44, 12113–12119.

(74) Lopes, D. H. J.; Meister, A.; Gohlke, A.; Hauser, A.; Blume, A.; Winter, R. Mechanism of Islet Amyloid Polypeptide Fibrillation at Lipid Interfaces Studied by Infrared Reflection Absorption Spectroscopy. Biophys. J. 2007, 93, 3132–3141.

(75) Lee, C. C.; Sun, Y.; Huang, H. W. How Type II Diabetes-Related Islet Amyloid Polypeptide Damages Lipid Bilayers. Biophys. J. 2012, 102, 1059–1068.

(76) Vorobieva, A. A.; White, P.; Liang, B.; Horne, J. E.; Bera, A. K.; Chow, C. M.; Gerben, S.; Marx, S.; Kang, A.; Stiving, A. Q.; et al. De Novo Design of Transmembrane b Barrels. Science 2021, 371.

(77) Mirecka, E. A.; Feuerstein, S.; Gremer, L.; Schröder, G. F.; Stoldt, M.; Willbold, D.; Hoyer, W. β-Hairpin of Islet Amyloid Polypeptide Bound to an Aggregation Inhibitor. Sci. Rep. 2016, 6, 1–8.

(78) Dupuis, N. F.; Wu, C.; Shea, J.; Bowers, M. T. Human Islet Amyloid Polypeptide Monomers Form Ordered β-Hairpins: A Possible Direct Amyloidogenic Precursor. J. Am. Chem. Soc. 2009, 131, 18283–18292.

(79) Reddy, A. S.; Wang, L.; Singh, S.; Ling, Y. L.; Buchanan, L.; Zanni, M. T.; Skinner, J. L.; De Pablo, J. J. Stable and Metastable States of Human Amylin in Solution. Biophys. J. 2010, 99, 2208–2216.

(80) Wu, C.; Shea, J. E. Structural Similarities and Differences between Amyloidogenic and Non-Amyloidogenic Islet Amyloid Polypeptide (IAPP) Sequences and Implications for the Dual Physiological and Pathological Activities of These Peptides. PLoS Comput. Biol. 2013, 9, 1–12.

(81) Raleigh, D.; Zhang, X.; Hastoy, B.; Clark, A. The β-Cell Assassin: IAPP Cytotoxicity. J. Mol. Endocrinol. 2017, 59, R121–R140.

(82) Liu, C.; Zhao, M.; Jiang, L.; Cheng, P. N.; Park, J.; Sawaya, M. R.; Pensalfini, A.; Gou, D.; Berk, A. J.; Glabe, C. G.; et al. Out-of-Register β-Sheets Suggest a Pathway to Toxic Amyloid Aggregates. Proc. Natl. Acad. Sci. U. S. A. 2012, 109, 20913–20918.

(83) Qiao, Q.; Wei, G.; Yao, D.; Song, Z. Formation of α-Helical and β-Sheet Structures in Membrane-Bound Human IAPP Monomer and the Resulting Membrane Deformation. Phys. Chem. Chem. Phys. 2019, 21, 20239–20251.

(84) Zhao, J.; Luo, Y.; Jang, H.; Yu, X.; Wei, G.; Nussinov, R.; Zheng, J. Probing Ion Channel Activity of Human Islet Amyloid Polypeptide (Amylin). Biochim. Biophys. Acta - Biomembr. 2012, 1818, 3121–3130.

(85) Zhao, J.; Hu, R.; Sciacca, M. F. M.; Brender, J. R.; Chen, H.; Ramamoorthy, A.; Zheng, J. Non-Selective Ion Channel Activity of Polymorphic Human Islet Amyloid Polypeptide (Amylin) Double Channels. Phys. Chem. Chem. Phys. 2014, 16, 2368–2377.

(86) Jang, H.; Arce, F. T.; Ramachandran, S.; Capone, R.; Lal, R.; Nussinov, R. Beta-Barrel Topology of Alzheimer’s Beta-Amyloid Ion Channels. J. Mol. Biol. 2010, 404, 917–934.

(87) Nguyen, P. H.; Campanera, J. M.; Ngo, S. T.; Loquet, A.; Derreumaux, P. Tetrameric Aβ40 and Aβ42 β-Barrel Structures by Extensive Atomistic Simulations. I. in a Bilayer Mimicking a Neuronal Membrane. J. Phys. Chem. B 2019, 123, 3643–3648.

(88) Chang, Z.; Luo, Y.; Zhang, Y.; Wei, G. Interactions of Aβ25-35 β-Barrel-like Oligomers with Anionic Lipid Bilayer and Resulting Membrane Leakage: An All-Atom Molecular Dynamics Study. J. Phys. Chem. B 2011, 115, 1165–1174.

(89) Knight, J. D.; Hebda, J. A.; Miranker, A. D. Conserved and Cooperative Assembly of Membrane-Bound α-Helical States of Islet Amyloid Polypeptide. Biochemistry 2006, 45, 9496–9508.

(90) Nath, A.; Miranker, A. D.; Rhoades, E. A Membrane-Bound Antiparallel Dimer of Rat Islet Amyloid Polypeptide. Angew. Chemie - Int. Ed. 2011, 50, 10859–10862.

(91) Harroun, T. A.; Bradshaw, J. P.; Ashley, R. H. Inhibitors Can Arrest the Membrane Activity of Human Islet Amyloid Polypeptide Independently of Amyloid Formation. FEBS Lett. 2001, 507, 200–204.

(92) Xiao, D.; Fu, L.; Liu, J.; Batista, V. S.; Yan, E. C. Y. Amphiphilic Adsorption of Human Islet Amyloid Polypeptide Aggregates to Lipid/Aqueous Interfaces. J. Mol. Biol. 2012, 421, 537–547.

(93) Saafi, E. L.; Konarkowska, B.; Zhang, S.; Kistler, J.; Cooper, G. J. S. Ultrastructural Evidence That Apoptosis Is the Mechanism by Which Human Amylin Evokes Death in RINm5F Pancreatic Islet β-Cells. Cell Biol. Int. 2001, 25, 339–350.

(94) Ritzel, R. A.; Butler, P. C. Islet Amyloid Polypeptide – Induced Apoptosis. Diabetes 2003, 52, 1701–1708.

(95) Brender, J. R.; Hartman, K.; Reid, K. R.; Kennedy, R. T.; Ramamoorthy, A. A Single Mutation in the Nonamyloidogenic Region of Islet Amyloid Polypeptide Greatly Reduces Toxicity. Biochemistry 2008, 47, 12680–12688.

(96) Magzoub, M.; Miranker, A. D. Concentration-dependent Transitions Govern the Subcellular Localization of Islet Amyloid Polypeptide. FASEB J. 2012, 26, 1228–1238.

(97) Akter, R.; Bower, R. L.; Abedini, A.; Schmidt, A. M.; Hay, D. L.; Raleigh, D. P. Amyloidogenicity, Cytotoxicity, and Receptor Activity of Bovine Amylin: Implications for Xenobiotic Transplantation and the Design of Nontoxic Amylin Variants. ACS Chem. Biol. 2018, 13, 2747–2757.

(98) Akter, R.; Abedini, A.; Ridgway, Z.; Zhang, X.; Kleinberg, J.; Schmidt, A. M.; Raleigh, D. P. Evolutionary Adaptation and Amyloid Formation: Does the Reduced Amyloidogenicity and Cytotoxicity of Ursine Amylin Contribute to the Metabolic Adaption of Bears and Polar Bears? Isr. J. Chem. 2017, 57, 750–761.

(99) Last, N. B.; Rhoades, E.; Miranker, A. D. Islet Amyloid Polypeptide Demonstrates a Persistent Capacity to Disrupt Membrane Integrity. Proc. Natl. Acad. Sci. U. S. A. 2011, 108, 9460–9465.

(100) Liu, G.; Prabhakar, A.; Aucoin, D.; Simon, M.; Sparks, S.; Robbins, K. J.; Sheen, A.; Petty, S. A.; Lazo, N. D. Mechanistic Studies of Peptide Self-Assembly: Transient α-Helices to Stable β-Sheets. J. Am. Chem. Soc. 2010, 132, 18223–18232.

(101) Godin, E.; Nguyen, P. T.; Zottig, X.; Bourgault, S. Identification of a Hinge Residue Controlling Islet Amyloid Polypeptide Self-Assembly and Cytotoxicity. J. Biol. Chem. 2019, 294, 8452–8463.

(102) Tian, Y.; Liang, R.; Kumar, A.; Szwedziak, P.; Viles, J. H. 3D-Visualization of Amyloid-β Oligomer Interactions with Lipid Membranes by Cryo-Electron Tomography. Chem. Sci. 2021, 12, 6896–6907.

(103) Sakagashira, S.; Hiddinga, H. J.; Tateishi, K.; Sanke, T.; Hanabusa, T.; Nanjo, K.; Eberhardt, N. L. S20G Mutant Amylin Exhibits Increased in Vitro Amyloidogenicity and Increased Intracellular Cytotoxicity Compared to Wild-Type Amylin. Am. J. Pathol. 2000, 157, 2101–2109.

(104) Ma, Z.; Westerrnark, G. T.; Sakagashira, S.; Sanke, T.; Gustavsson, A.; Sakarnoto, H.; Engstrorn, U.; Nanjo, K.; Westermark, P. Enhanced in Vitro Production of Amyloid-like Fibrils from Mutant (S20G) Islet Amyloid Polypeptide. Amyloid 2001, 8, 242–249.

(105) Cao, P.; Tu, L. H.; Abedini, A.; Levsh, O.; Akter, R.; Patsalo, V.; Schmidt, A. M.; Raleigh, D. P. Sensitivity of Amyloid Formation by Human Islet Amyloid Polypeptide to Mutations at Residue 20. J. Mol. Biol. 2012, 421, 282–295.

(106) Meier, D. T.; Entrup, L.; Templin, A. T.; Hogan, M. F.; Mellati, M.; Zraika, S.; Hull, R. L.; Kahn, S. E. The S20G Substitution in HIAPP Is More Amyloidogenic and Cytotoxic than Wild-Type HIAPP in Mouse Islets. Diabetologia 2016, 59, 2166–2171.

(107) Bucciantini, M.; Giannoni, E.; Chiti, F.; Baroni, F.; Taddei, N.; Ramponi, G.; Dobson, C. M.; Stefani, M. Inherent Toxicity of Aggregates Implies a Common Mechanism for Protein Misfolding Diseases. Nature 2002, 416, 507–511.

